# A heterohexameric protein consisting of six linked single-domain antibodies is highly protective for BoNT/A, BoNT/B and BoNT/E exposures

**DOI:** 10.1101/2021.03.02.433625

**Authors:** Jean Mukherjee, Jacqueline M. Tremblay, Michelle Debatis, Alexa Foss, Junya Awata, Charles B. Shoemaker

## Abstract

Botulinum neurotoxin (BoNT) serotypes A, B and E cause the vast majority of human botulism cases and pose the greatest bioterrorism threats. We previously identified multiple camelid single-domain antibodies (VHHs) that each neutralize BoNT/A, BoNT/B or BoNT/E. We also demonstrated that heterodimers of linked toxin-neutralizing VHHs are much more potent than VHH monomer pools in preventing BoNT intoxication. In this study, we expressed two different heterohexamer proteins (VNA1-ABE and VNA2-ABE) of ~100 kDa secreted from mammalian host cells, each containing the same six linked anti-BoNT VHH components ordered in two different combinations. Each heterohexamer contained two VHHs that neutralize BoNT/A, BoNT/B or BoNT/E. Both heterohexameric antitoxins displayed similar strong binding properties for the three targeted BoNT serotypes by ELISA. One ug of each heterohexameric antitoxin fully protected groups of mice co-administered with 100 LD_50_ of BoNT/A, BoNT/B or BoNT/E, or a pool containing 100 LD_50_ of each of the three toxins. The results demonstrate that long chains of at least six different linked VHHs can be expressed such that all component VHHs in the multimer retain their target binding activities. These findings make more feasible the development of a BoNT antitoxin product consisting of a small pool of proteins that, in combination, neutralize all known BoNT serotypes and subtypes.

**Key Contribution:** Heteromultimeric proteins consisting of six linked, VHH antibodies, and including VHHs that neutralize BoNT/A, BoNT/B and BoNT/E, retain high potency to protect mice challenged with high doses of all three of these BoNT serotypes.

## 1. Introduction

Protein toxins are important virulence factors in many important diseases and can also cause harm through accidental exposures or deliberate bioterror incidents. Specific antitoxin antibodies (Abs) are the primary treatment modality for toxin exposures. Traditionally, antitoxins are either elicited in patients through vaccination or passively administered as antiserum or purified Abs. Immunization is not practical for most toxin threats and traditional antitoxin treatments are often expensive, difficult to access and may pose safety concerns. Thus, new therapeutic options that mitigate these issues are needed.

One alternative to conventional serum-based antitoxins are single-domain Abs (sdAbs) generated from camelid [1] or shark [2] heavy-chain only Abs (HcAbs). These simple Abs are the V_H_ region of HcAbs and are often called nanobodies or VHHs. Advantages of VHHs over conventional Abs are that VHHs are small (~14 kDa) proteins, more stable to pH and temperature extremes, and can be produced inexpensively in microbial hosts [3–5]. In addition, VHHs can be expressed as multimeric proteins which impart improved antitoxin potencies [6–8] and permit the targeting of multiple toxins with a single protein agent [9–13].

VHH antitoxins also have some disadvantages as compared to conventional antibodies. One disadvantage is that VHHs are much more rapidly cleared from the circulation than conventional antibodies. Since antitoxins are generally administered in a single post-exposure treatment, serum persistence is not important if the treatment adequately neutralizes the toxin that had already entered the system. For situations in which longer serum persistence is needed, the antitoxin VHHs can be fused to a second VHH or peptide having high affinity for a major serum protein such as albumin [7, 14]. Alternatively, VHHs are easily amenable to genetic delivery and can be expressed for extended times in patients by gene therapy with viral vectors [7] or delivered by encapsulated synthetic mRNA [15]. A second potential disadvantage of VHHs is that they lack the Fc effector functions of conventional antibodies, which results in the inability of VHHs to promote clearance of the toxin from the circulation. Treatment with polyclonal antisera results in the decoration of the toxin with multiple Ab Fc domains, and it is known that decoration by at least three Fc domains leads to rapid serum clearance [16]. Promotion of toxin serum clearance, though, should not be necessary if the antitoxin agents adequately neutralize the toxin until natural processes clear the toxin. Indeed, HBAT (heptavalent botulinum antitoxin, Cangene Corporation), which is the current treatment for exposure to *Clostridium botulinum* neurotoxins (BoNT), consists of polyclonal Fab domains purified from horse antiserum. Thus HBAT, like VHHs, lacks Fc domains, resulting in rapid product clearance from serum [17] and an inability to promote Fc receptor-mediated toxin clearance. For situations in which an Fc domain is necessary for a VHH-based antitoxin product to be fully effective, it is possible to produce the VHH agents as fusions to Fc domains [18] or to co-administer an anti-tag effector Ab which binds to peptide tags engineered into the VHH products [8, 11, 19] and thus provide Fc functions.

Most protein toxin threats to humans derive from pathogens that exist in nature as families of related microorganisms that produce natural variant forms of one or more toxins. BoNTs are an extreme example of this challenge as at least seven distinct BoNT serotypes have been identified [20], most of which can be found with diverse natural subtype variations. Antitoxin products must be capable of broadly neutralizing all potential variants of the target toxin. The use of polyclonal antitoxin sera to treat toxin exposures generally provides broad protection from toxin variants since it consists of a diverse pool of antibodies recognizing many different epitopes, at least a few of which are likely to be effective against each of the variants. Efforts to replace polyclonal antitoxin products with clonal antibody products such as VHHs must also be capable of broadly neutralizing natural variants. One solution that is being developed for BoNTs is to create pools of three mAbs for each serotype in which each mAb has been carefully selected for the ability to broadly neutralize all known natural variant subtypes for the serotype and each binds their target at non-overlapping epitopes. With this design, each mAb pool should neutralize virtually all natural subtype variants and decorate each toxin with three mAb Fc domains so as to also promote rapid toxin clearance from the serum [21–26].

VHH antitoxins may be engineered to become capable of broad neutralization of natural variants through production of heteromultimers containing a series of different linked VHHs, each selected for the ability to broadly neutralize a subset of natural toxin variants. Even when both VHH components of a heterodimer have relatively poor neutralizing activity on a toxin as monomers, the linked VHHs can acquire high potency [11]. Prior studies have demonstrated that VHH heterotetramers with four different toxin neutralizing VHHs can be produced and that each VHH component appears to fold properly during expression and remain functional [9]. These heteromultimeric proteins are called VHH-based neutralizing agents or VNAs. In this study, we tested the feasibility of expressing a VHH heterohexameric VNA in which two VHH components each recognize one of three different BoNT serotypes; BoNT/A, BoNT/B and BoNT/E. The heterohexamer components are organized in two different formats and both proteins tested for the ability to bind to each of the three toxins and to protect mice from lethal intoxication by the toxins.

## 2. Results

### 2.1. Expression of VHH heterohexamer proteins

Two synthetic genes, each encoding six linked VHH components, were designed and prepared. Both heterohexameric VNAs contained the same six VHHs arranged in a different order (**Figure 1A**). The six VHHs selected as components consisted of two VHHs that were each known to potently neutralize one of three different toxins; BoNT/A, BoNT/B or BoNT/E. The two component VHHs that neutralize BoNT/A are ciA-H7 (A1) and ciA-B5 (A2) [19]; for BoNT/B they are JLI-G10 (B1) and JLK-G12 (B2) [27]; and for BoNT/E they are, JLE-G6 (E1) and JLE-E9 (E2) [28]. All VHHs in the VNAs recognized non-overlapping neutralizing epitopes on their different toxin targets. The six VHHs were linked in two different combinations as shown in **Figure 1A**, and the complete monomer and heterohexamer sequences are provided in **Figure S1A and S1B**. The first heterohexamer construction, VNA1-ABE contains the pairs of VHHs recognizing each BoNT serotype linked together such that the order from amino to carboxyl end would be A1/A2/B1/B2/E1/E2. In VNA1-ABE, all VHHs were linked together by flexible spacers that match the spacers of VHH heterodimers for each BoNT serotype that had been successfully tested as antitoxin in mice [19, 27, 28]. The second VHH heterohexameric VNA construction, VNA2-ABE, contains the pairs of VHHs recognizing each serotype separated from one another by two VHHs targeting a different BoNT serotype, with the order A1/B1/E1/A2/B2/E2. In VNA2-ABE, we employed shorter, 5 amino acid GGGGS spacers to minimize overall protein size.

**Figure 1.**
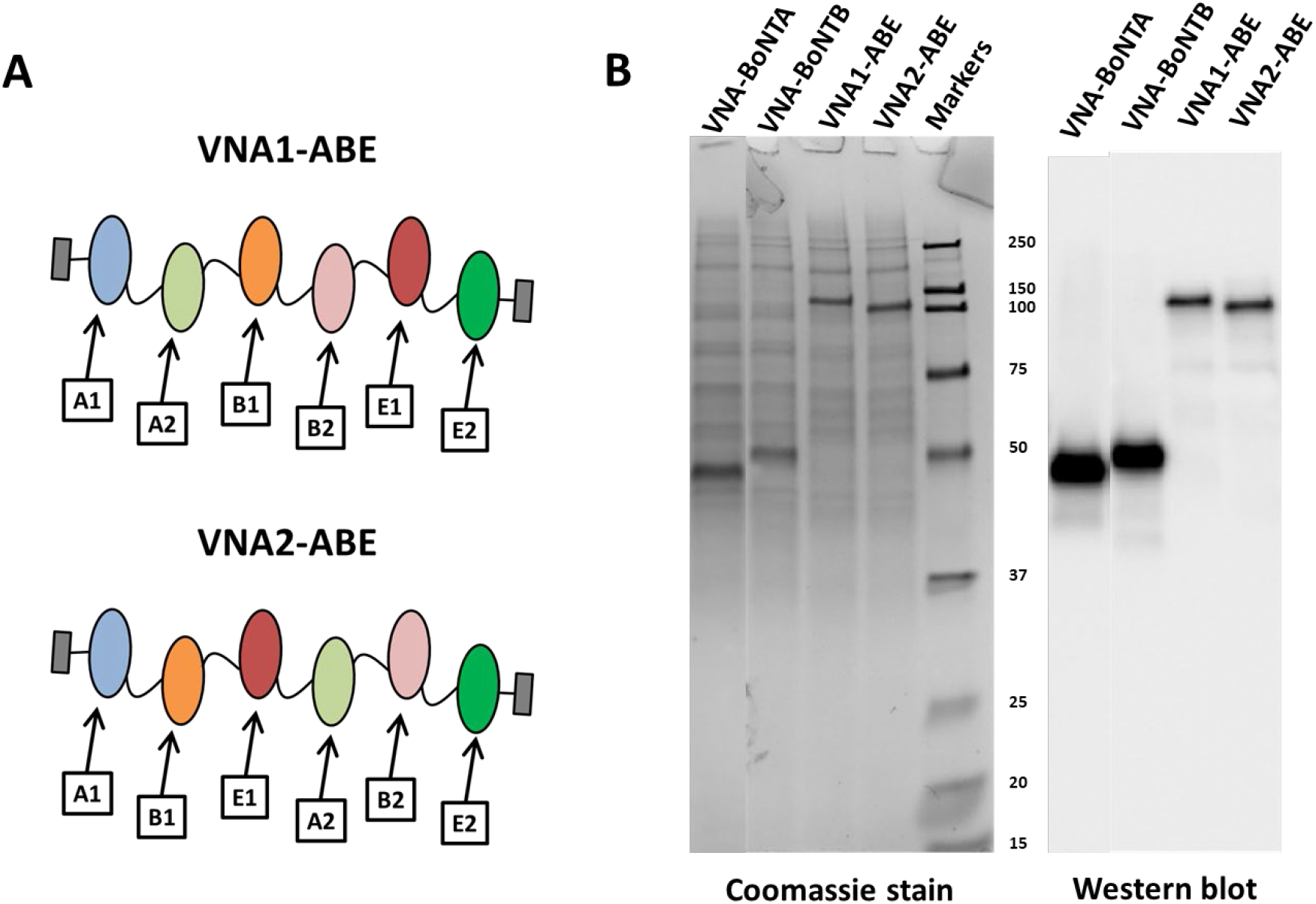
Design and expression of heterohexameric VNA antitoxins. A. Cartoon image showing the VHH component structures of VNA1-ABE and VNA2-ABE. The sequences of the six component VHHs: A1, A2, B1, B2, E1, E2 are shown in **Figure S1A**. The full amino acid sequences of the two VNA heterohexamers are shown in **Figure S1B**. B. Conditioned medium was obtained from the CHO cells transfected with vectors encoding the four different indicated VNAs and samples were loaded onto SDS-PAGE for Coomassie staining (10 μl) or western blotting (1 μl). Western blots were probed with HRP/anti-E-tag, molecular weight marker sizes are shown in kb.

The DNAs encoding VNA1-ABE and VNA2-ABE were ligated into a mammalian cell expression vector, pSecB, under the control of the CMV promoter and fused to the human Igκ leader sequence. Similar pSecB vectors were prepared containing heterodimers of BoNT/A neutralizing VHHs A1/A2 (VNA-BoNTA) and BoNT/B-neutralizing VHHs, B1/B2 (VNA-BoNTB). The plasmids were transfected into CHO cells grown in serum-free medium and the conditioned medium collected after 3-4 days. The conditioned medium was characterized by SDS-PAGE followed by protein staining or Western blotting (**Figure 1B**). The products of expected size were visible by protein staining of conditioned medium and their identities confirmed by the blots. The blots also revealed that there were no detectable truncated VNA products of the heterohexamers in the conditioned medium. The expression levels for both heterohexamer proteins were estimated to be approximately 10 μg/ml (~100 nM) based on comparisons of the staining intensity against protein standards. Similar protein expression levels were obtained in conditioned media from cells transfected with the two heterodimer VNA vectors (~10 μg/ml (~300 nM)).

### 2.2. VHH heterohexamers recognize each of their three BoNT targets

Mammalian cell produced recombinant VNA1-ABE and VNA2-ABE VHH hexamer proteins were assessed for their ability to bind to each of the three BoNT serotypes to which the monomer components recognize. ELISAs were performed to assess VNA binding to catalytically inactive forms of BoNT/A, BoNT/B or BoNT/E holotoxins [29, 30]. As shown in **Figure 2**, each of the two hexameric VNAs recognized all three toxins with approximately equal apparent affinities based on their similar EC_50_ between ~0.5 and 2 nM. The VNA-BoNTA and VNA-BoNTB heterodimers, also expressed in mammalian cells, possessed similar apparent affinities (EC_50_s) to the two heterohexamers tested on ciBoNTA and ciBoNTB. The VNA-BoNTE heterodimer, which was expressed in *E. coli*, had an EC_50_ similar to the two heterohexamers when tested on ciBoNTE. The EC_50_ values reflecting apparent affinity likely under-represent the actual affinity of these agents for their native BoNT targets because we employed ELISAs with plastic-coated ciBoNTs so as to be consistent across the three BoNT serotypes. Other studies have shown that VHHs typically recognize the more conformationally-native antibody-captured BoNT targets with higher apparent affinity than the plastic-coated BoNTs [28]. Nevertheless, the results demonstrate that the apparent affinity of the two heterohexamers for their three different BoNT target serotype targets did not detectably differ from that of comparable VHH heterodimers, despite their expression as a single chain of six linked VHH components.

**Figure 2.**
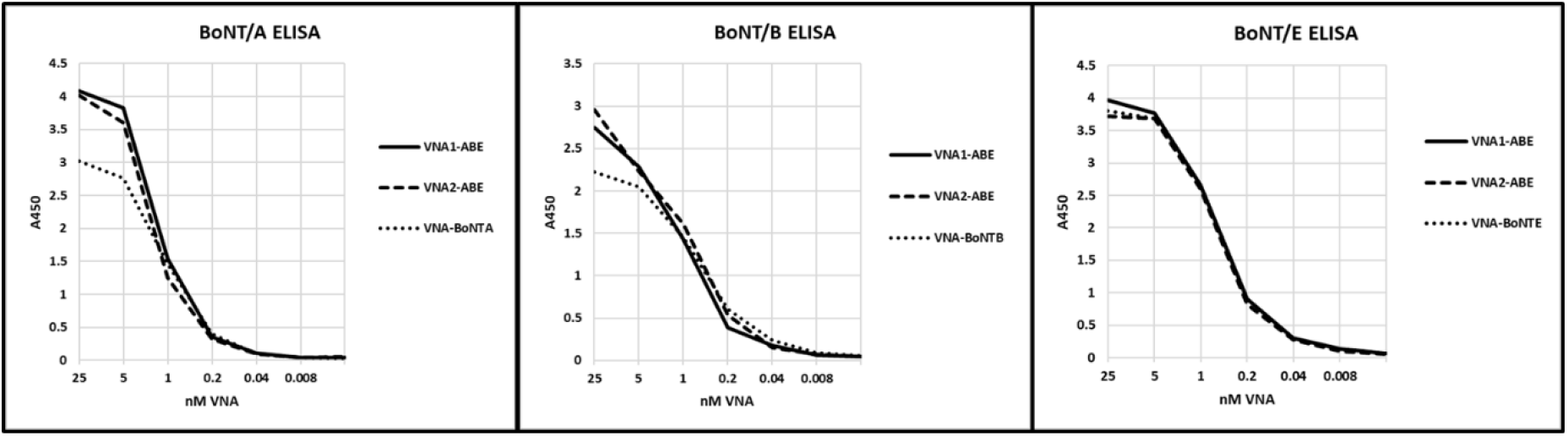
Dilution ELISAs of heterohexamer and heterodimer VNAs on different BoNT serotypes. ELISAs were performed by coating 1 μg/ml of the target BoNT (ciBoNTA, ciBoNTB or ciBoNTE) onto Costar tissue culture 96-well plates. All VNAs were diluted to 25 nM and then subjected to 1:5 serial dilutions. VHH binding was detected with HRP/anti-E-tag reagent.

### 2.3. VHH heterohexamers protect mice from intoxication by their three BoNT targets

Several in vivo mouse studies were performed to assess the antitoxin efficacies of the two heterohexamers. **Figure 3** shows survival plots for groups of five mice challenged with 100 LD_50_ of each BoNT serotype, or a pool containing 100 LD_50_ of each of the three BoNT serotypes. All control mice died <4 hours following toxin exposure while all mice given 10 pmoles (1 μg) VNA1-ABE or VNA2-ABE survived challenge with either the individual or combined toxin challenge. All treated mice developed transient mild-moderate signs of botulism and recovered within a week. Previous studies have shown that use of a pool of two unlinked BoNT/A-neutralizing VHHs, ciA-H7 (A1) and ciA-B5 (A2) are much less potent as antitoxins than a heterodimer containing the same VHH components and unable to protect mice challenged with 100 LD_50_ of BoNT/A [19]. The two BoNT/E-neutralizing monomer component, JLE-G6 (E1) or JLE-E9 (E2), were unable to protect mice from as low as 3 LD_50_ of BoNT/E [28]. These results strongly suggest that the VHH monomer components of the heterohexamer remain linked together within the treated animals.

**Figure 3.**
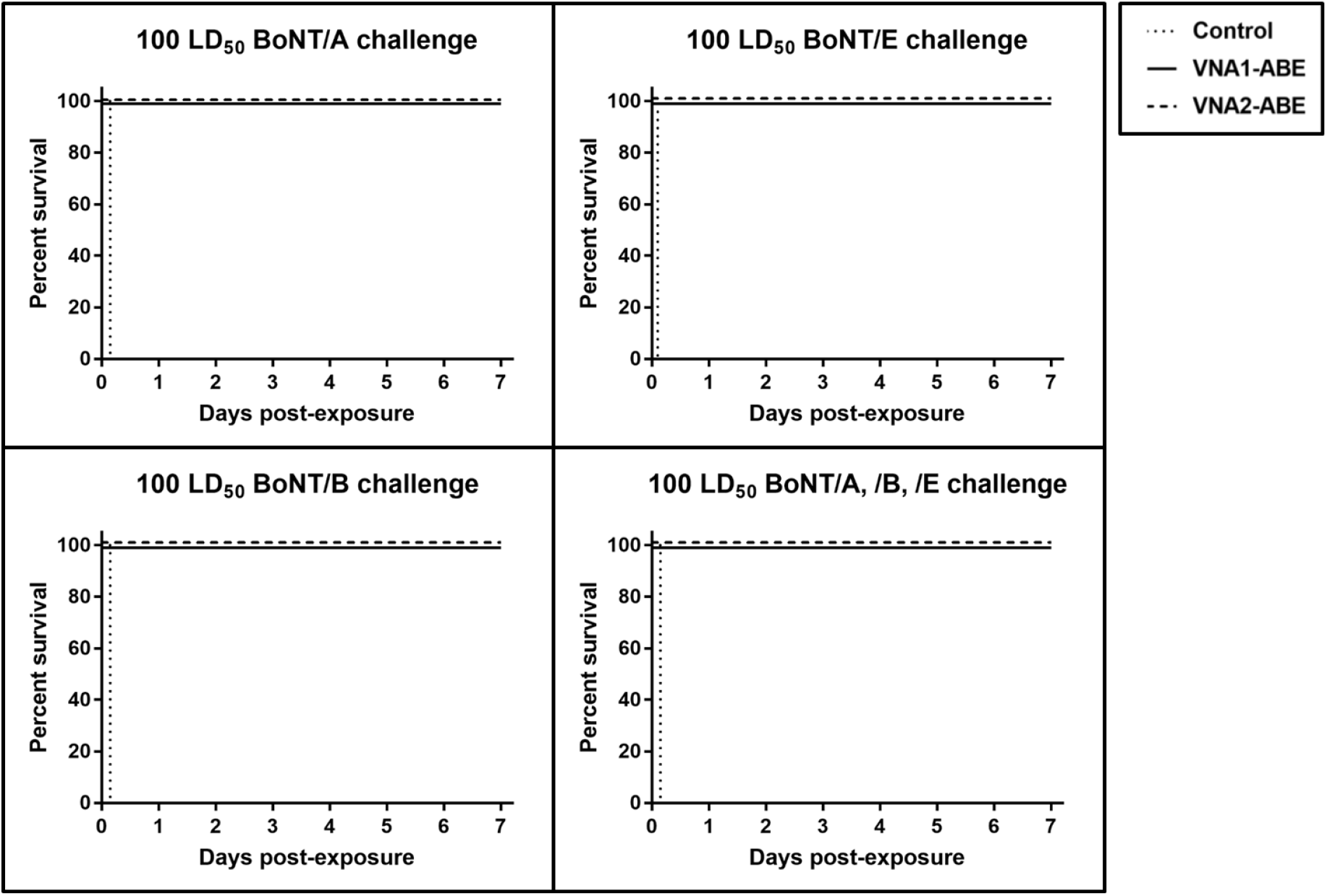
Heterohexamer VNAs protect mice from 100 LD_50_ doses of BoNT/A, BoNT/B and BoNT/E. Groups of five mice were co-administered 1 μg (~10 pmoles) of VNA1-ABE or VNA2-ABE and 100 LD_50_ of BoNT/A, BoNT/B or BoNT/E, or a combination of all three toxins, as indicated. Mice were then monitored for survival and signs of botulism for a week. The survival of mice is plotted as a function of time post-exposure. All VNA-treated groups had 100% survival, though showing mild signs of botulism, and these outcomes were significantly different from untreated controls (p<0.001 by log-rank Mantel-Cox test).

Additional in vivo studies with the heterohexamer proteins compared the survival of mice challenged with 500 LD_50_ of BoNT/B that were co-administered with heterohexamers VNA1-ABE, VNA2-ABE or the heterodimer, VNA-BoNTB. As shown in **Figure 4A**, control mice died within two hours while all mice treated with the three VNAs survived, showing only mild signs of botulism before recovering. Finally, groups of mice were challenged with 10,000 LD_50_ of BoNT/A co-administered with 10 pmoles of heterohexamers VNA1-ABE, VNA2-ABE or 10 pmoles of heterodimer VNA-BoNTA (**Figure 4B**). Control mice died in less than an hour while mice treated with the VNAs survived about for one or two days. These results suggest there may be small differences in potency between VNA1-ABE and VNA2-ABE revealed at the highest challenge dose, although this would need to be confirmed by additional studies.

**Figure 4.**
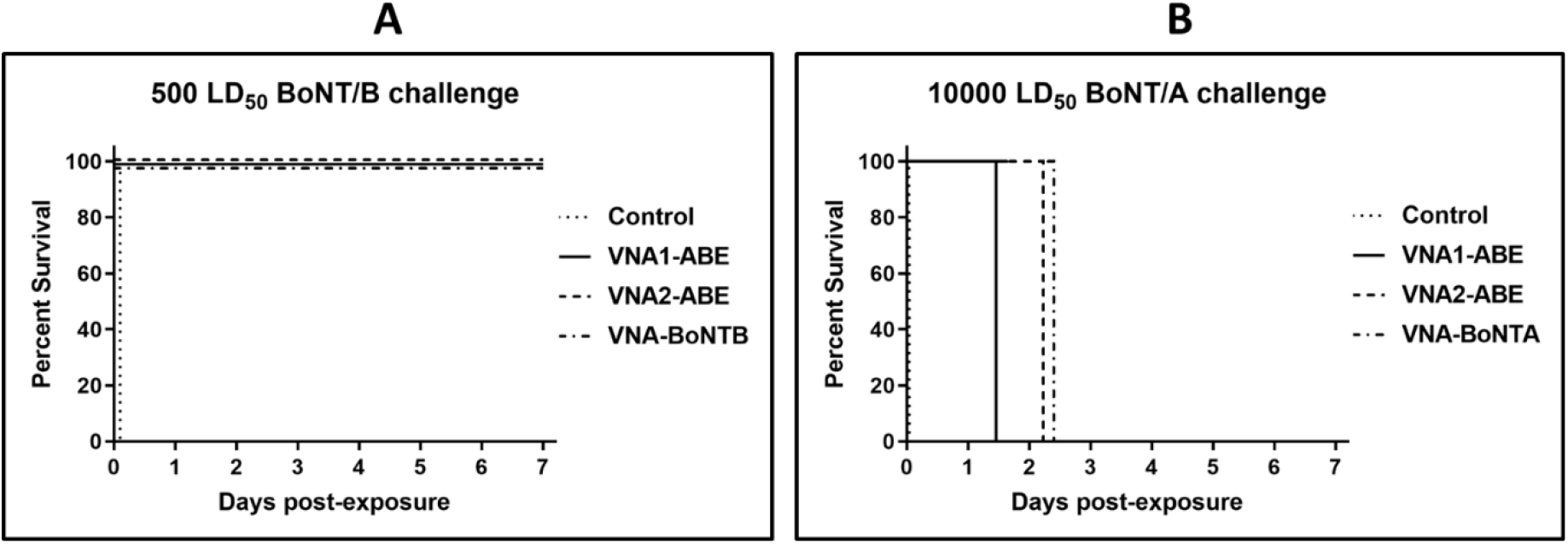
Heterohexamer VNAs protect mice from 500 LD_50_ doses of BoNT/B and delay death from 10,000 LD_50_ doses of BoNT/A. A. Groups of five mice were co-administered 2 μg of VNA1-ABE, VNA2-ABE (20 pmoles), or 1 ug of VNA-BoNTB (20 pmoles) and 500 LD_50_ of BoNT/B. Mice were then monitored for survival and signs of botulism for a week. The survival of mice is plotted as a function of time post-exposure. All VNA-treated groups had 100% survival, significantly different from untreated controls (p<0.001 by log-rank Mantel-Cox test). B. Groups of five mice were co-administered 1 μg of VNA1-ABE, VNA2-ABE (10 pmoles), or 2 ug of VNA-BoNTA (40 pmoles) and 10,000 LD_50_ of BoNT/A. Mice were then monitored for survival and signs of botulism. The survival of mice is plotted as a function of time post-exposure. Treated groups differed significantly from untreated controls (p<0.01 by log-rank Mantel-Cox test).

## 3. Discussion

VHHs have proven to be excellent components of BoNT antitoxin agents when produced as linked multimers [19, 27, 28]. The ability to express VHHs as multimers makes possible the ability to develop therapeutic antitoxins targeting multiple different toxins, such as *C. difficile* toxins TcdA and TcdB [9, 13], anthrax EF and LF [12] or Shiga toxins Stx1 and Stx2 [10], with a single protein. Here we extend these findings to show that a single heterohexameric protein containing six different linked VHH components makes possible antitoxin protection from three different BoNT serotypes. Since the heterohexamers retained the same apparent affinity and antitoxin potencies as the heterodimers for each serotype, these results also indicate that all component VHHs retain their full binding and neutralizing functions. This implies that even the VHH components located proximal to the carboxyl end of the growing chain of six linked VHHs continue to fold into independent functioning domains during translation and secretion.

Two different heterohexamer constructions were produced; one in which VHH pairs targeting the same toxin serotype were placed adjacent to one another, and the second construction in which the two VHHs targeting the same serotype were separated by two VHHs targeting a different serotype (Figure 1A). We were unable to identify any clear differences between the two different constructions in terms of their apparent affinity for their toxin targets or their ability to protect mice from intoxication by the different BoNT serotypes. A previous study has indicated that heterodimeric VNAs in which both VHH components are sterically capable of binding simultaneously to the toxin are more potent antitoxins [27]. Our results show that separating the VHH components with several bulky VHH components did not demonstrably improve the potency and thus suggest that the VHHs in this heterodimer did not permit simultaneous binding of both VHH components to the same toxin molecule despite the presence of two intervening linked VHHs. Furthermore, the fact that both VNA heterohexamers displayed the same antitoxin potencies as the individual heterodimers for each serotype strongly suggests that all six of the individual VHH components retained their full toxin binding functions. This is most well demonstrated for BoNT/E as both anti-BoNT/E VHH components are present at the carboxyl terminus of the heterohexamer, the last components synthesized during translation, and yet the heterohexamer retained the same anti-BoNT/E binding affinity by ELISA and the same antitoxin potency in vivo as the VNA/BoNTE heterodimer.

Antitoxins for botulism, or any toxin, should possess the ability to provide protection from most if not all natural variants of the toxin to which they may be exposed. Use of antitoxins that derive from polyclonal antibodies generally offer protection to natural variants because they consist of many different antibodies recognizing different toxin epitopes, thus typically having a subset of cross-specific antibodies that recognize each of the variants. But natural variants create a challenge for antitoxin products consisting of clonal antibody components. For example, monoclonal antibodies (mAbs) must be identified that broadly recognize all known natural variants or, alternatively, pools of mAbs must be produced that, in combination, provide broad variant protection. The ability to produce long chains of VHH components, up to at least six different VHHs as shown here, offers an alternative solution as it permits the incorporation of numerous VHHs into one or a few proteins designed to neutralize virtually all of the natural diversity of the toxin target. While such an idealized goal is particularly challenging for a botulism antitoxin, we suggest that it should be possible to produce a broad BoNT antitoxin consisting of perhaps only three or four heteromultimeric proteins. This would be accomplished by selecting the appropriate BoNT-neutralizing VHH components such that all naturally occurring BoNT serotypes and subtypes are targeted for neutralization by at least two of the VHH components.

One attractive option for a next-generation BoNT antitoxin might be nanoparticle-encapsulated RNA encoding three or four heteromultimer antitoxin proteins that together possess good potency to all known BoNT serotypes and subtypes in nature. We have reported the successful use of RNA nanoparticles to elicit high levels of expression of antitoxin VNAs in mice [15] as a novel form of passive immunization. Mice were shown to express VNA serum levels that exceeded 100 ug/ml at day 1 post treatment and fully protected mice from lethal challenges of either BoNT/A or BoNT/B in mice. Prior studies have shown that only about 10 ng of VNA is sufficient to protect mice from 10 LD_50_ of BoNT/A [7] which is 10,000-fold below the levels produced in the RNA treated mice. More importantly, the serum VNAs were easily detectable in mice within 2 hours post-treatment and the post-intoxication treatment window was indistinguishable between mice treated with RNA nanoparticles and those given intravenous protein VNA [15].

## 4. Conclusions

We have demonstrated that VHH heteromultimers of six VHHs can be efficiently expressed and secreted in mammalian cells with the retention of the full function of each VHH component. We also demonstrate that a single VHH heterohexamer can be produced that effectively protects animals from intoxication by three different toxins, in this case BoNT/A, BoNT/B and BoNT/E. These findings suggest the feasibility of developing a broad specificity BoNT antitoxin capable of treating recent exposures from virtually any natural BoNT serotype or subtype with a small pool of VHH multimers. Furthermore, this work highlights the great potential of VHH antibodies to retain high their individual functionality even when expressed as chains of linked VHHs containing at least six VHH components.

## 5. Materials and Methods

### 5.1. Ethics statement

All in vivo studies were performed within the guidelines established by the Guide for the Care and Use of Laboratory Animals of the National Institutes of Health and were approved by the Tufts University Institutional Animal Care and Use Committee (IACUC) and conducted under Protocol G2016-74.

### 5.2. Toxins

BoNT/A, BoNT/B and BoNT/E complex preparations (subtype 1 for each toxin, all from Metabiologics Inc.) were used for in vivo assays. Prior to use, BoNT/E was subjected to trypsin activation. Briefly, 25 ul 1.2 mg/ml trypsin (EMD #650211) in HEPES (Sigma #H0887), 50 ul 1 mg/ml Complexed Botulinum Toxin E and 100 ul sterile PBS were incubated 30 min. at 37°C. Following incubation, 25 ul 2.5 mg/ml trypsin inhibitor (Sigma #T6522) was added and the mixture further incubated 15 min. at room temperature. The resulting solution containing 0.25 ug/ul activated BoNT/E was stored at −80°C until use. BoNT/A, BoNT/B and the activated BoNT/E were carefully titrated in vivo to determine the LD50 prior to use in assays performed for VHH evaluation. All procedures utilizing BoNT were performed within a CDC-registered Select Agent laboratory.

### 5.3. VHH and VNA expression

VHHs and some VHH heterodimeric VNAs were expressed in *E. coli* as previously reported. For expression of most VNAs, a CHO cell host was employed. Coding DNA encoding the VNAs was synthesized by Genscript with restriction sites compatible with insertion in frame into a mammalian expression vector based on pSecTag2. The sequences of encoded heterohexamer proteins are shown in Supplemental Figure S1. Similar expression vectors were engineered for expression of heterodimeric VNAs encoding VHHs A1/A2 (VNA-BoNTA) or B1/B2 (VNA-BoNTB). Expression of the heterodimeric VNA-BoNTE containing E1/E2 is described as Trx/E/JLE-G6/JLE-E9/E in Tremblay et al [28] and was produced and purified from *E. coli*. VNAs contained E-tags tags for detection. The mammalian expression plasmids were transfected into CHO-S cells in suspension cultures of serum-free FreeStyle™ CHO cell medium. Cells were transfected using FreeStyle™ MAX reagent (Invitrogen) according to the manufacturer’s recommendations and incubated under orbital shaking for three days. Conditioned medium was harvested under sterile conditions and stored at 4°C. VHH and VNA expression levels were estimated by comparison to protein standards using BioRad Image Lab software.

### 5.4. ELISAs

ELISAs were performed using Costar tissue culture 96 well plates to reduce the conformational deformation that occurs with conventional ELISA plastic [28]. Catalytically inactive BoNT proteins representing BoNT/A, BoNT/B and BoNT/E [29–31] (kindly provided by Dr. Robert Webb, USAMRICD) were coated at 4°C overnight at 1 μg/ml in PBS, then blocked for at least an hour at 37°C with 4% milk in PBS, 0.1% Tween. After washing, dilution ELISAs were initiated by diluting the VNAs to 25 nM and performing serial dilutions of 1:5. After incubation for one hour at 37°C, plates were washed and then incubated with 1:10,000 rabbit HRP/anti-E-tag (Bethyl) for one hour, washed, developed with TMB (Sigma) as recommended by manufacturer and read at A450.

### 5.5 Standard mouse toxin lethality assay

VNA1-ABE and VNA2-ABE were evaluated in vivo using the BoNT murine lethality assay. The lethality assays utilized adult, ~20 g female CD-1 mice (Charles River Labs) housed in standard shoebox cages at 5 mice/cage. Following arrival, mice were observed at least twice daily and cages containing corncob bedding and nestlets along with standard rodent chow within the cage rack and water ad libitum were changed twice weekly. Throughout each study, mice were also offered food on the cage bottom along with both nutritional (Napa Nectar; Lenderking #88-0001) and hydration (Boost; Clear H2O #72-04-5022) supplement gels. One day prior to initiation of each study, mice were weighed and sorted to reduce intergroup (n=5) weight variation. VNA-BoNT mixtures were co-administered IV via the lateral tail vein within a 200-250 μl volume. Each mouse within the respective group received 1 μg of VNA1-ABE (10 pmoles) or VNA2-ABE (10 pmoles) and 100 LD_50_ of BoNT/A1, BoNT/B1 or BoNT/E or 100 LD_50_ BoNT/A1, BoNT/B1 and BoNT/E. Following administration of BoNT, mice were observed a minimum of four times each day between 9 a.m. and midnight with additional checks throughout peak periods of mortality to ensure accurate documentation of time to death data. Time to death was defined as the time at which a mouse was found dead or was euthanized due to severe morbidity. At each check, a standard score sheet was utilized to objectively document the overall disposition and clinical symptoms exhibited by each mouse. Mice which were observed to be open-mouth breathing, lethargic, unresponsive or moribund were euthanized via carbon dioxide asphyxiation followed by cervical dislocation.

## Supplementary Materials

The following are available online at www.mdpi.com/xxx/s1, Figure S1: Sequences of heterohexamer VNAs and their component VHHs.

## Author Contributions

Conceptualization, C.B.S., methodology, J.M., J.M.T, M.D., A.F., J.A, formal analysis J.M., C.B.S., investigation, J.M., J.M.T, M.D., A.F., J.A, data curation, J.M., J.M.T, M.D., A.F, supervision, J.M., C.B.S., writing, J.M., C.B.S., funding acquisition, C.B.S. All authors have read and agreed to the published version of the manuscript.

## Acknowledgments

We gratefully acknowledge Karen Baldwin, Kwasi Ofori and Zachary Forbes for expert technical assistance with handling of alpacas, isolation of PBLs and performance of mouse LD50 assays which made these subsequent studies possible. We are also grateful to Dr. Patrick M. McNutt for help with some of the statistical analysis.

