## Supplemental Figure S1 for "A heterohexameric protein consisting of six linked single-domain antibodies is highly protective for BoNT/A, BoNT/B and BoNT/E exposures"

## A

| VHH | Name | Heterohexamer component VHH amino acid sequences |
|---|---|---|
| A1 | ciA-B5 | QLQLVESGGGLVHPGGSLRLSCAPSASLPSTPFNPFNNMVGWYRQAPGKQREMVASIGLRINYADSVKGRFTISRDNAKNTVDLQMDSLRPEDSATYYCHIEYTHYWGKGTLVTVSSEPKTPKPQ |
| A2 | ciA-H7 | QLQLVESGGGLVQVGGSLRLSCVVSGSDISGIAMGWYRQAPGKRREMVADIFSGGSTDYAGSVKGRFTISRDNAKKTSYLQMNNVKPEDTGVYYCRLYGSGDYWGQGTQVTVSSAHHSEDP |
| B1 | JLI-G10 | QLQLVESGGGLVQAGGSLRLSCAASILTYDLDYYYIGWVRQAPGKEREGVSCISSTDGATYYADSVKGRFTISRNNAKNTVYLQMNNLKPEDTAIYYCAAAPLAGRYCPASHEYGYWGQGTQVTVSSAHHSEDP |
| B2 | JLK-G12 | QLQLVESGGGLVQAGGSLRLSCAASEFRAEHFAVGWFRQAPGKEREGVSCVDASGDSTAYADSVKGRFTISRDNNKNVVYLQMDSLEPEDTGDYYCGASYFTVCAKSMRKIEYRYWGQGTQVTVSSEPKTPKP |
| E1 | JLE-G6 | QAQLQLVESGGGLVKPGGSLRLSCVVSGFTFDDYRMAWVRQAPGKELEWVSSIDSWSINTYYEDSVKGRFTISTDNAKNTLYLQMSSLKPEDTAVYYCAAEDRLGVPTINAHPSKYDYNYWGQGTQVTVSSEPKTPKP |
| E2 | JLE-E9 | QAQLQLVESGGGLVQAGGSLRLSCAASGRTFSSYSMGWFRQAPGKEREYVAAVNSNGDSTFYADSIKGRFTVSRDAAKNTVYLQMNSLKPEDTALYYCAAVYGRYTYQSPKSYEYWGQGTQVTVSSEPKTPKP |

## B

**VNA1-ABE**

**METDTLLLWVLLLWVPGSTGDAAQPARRARRTKLSGAPVPYPDPLEPRAAAGQGVQAQLQLVESGGGLVHPGGSLRLSCAPSASLPSTPFNPFNNMVGWYRQAPGKQREMVASIGLRINYADSVKGRFTISRDNAKNTVDLQMDSLRPEDSATYYCHIEYTHYWGKGTL VTVSSEPKTPKPQSGGGGSGGGGSGGGGSGGGGSGGGGSGQGVQAQLQLVESGGGLVQVGGSLRLSCVVSGSDISGIAMGWYRQAPGKRREMVADIFSGGSTDYAGSVKGRFTISRDNAKKTSYLQMNNVKPEDTGVYYCRLYGSGDYWGQGTQVTVSSAHHSEDP SGGGGQGVQAQLQLVESGGGLVQAGGSLRLSCAASILTYDLDYYYIGWVRQAPGKEREGVSCISSTDGATYYADSVKGRFTISRNNAKNTVYLQMNNLKPEDTAIYYCAAAPLAGRYCPASHEYGYWGQGTQVTVSSAHHSEDPSGGGGSGGGGSGGGGSGQGVQAQL QLVESGGGLVQAGGSLRLSCAASEFRAEHFAVGWFRQAPGKEREGVSCVDASGDSTAYADSVKGRFTISRDNNKNVVYLQMDSLEPEDTGDYYCGASYFTVCAKSMRKIEYRYWGQGTQVTVSSEPKTPKPQSGGGGQGVQAQLQLVESGGGLVKPGGSLRLSCVVSGF TFDDYRMAWVRQAPGKELEWVSSIDSWSINTYYEDSVKGRFTISTDNAKNTLYLQMSSLKPEDTAVYYCAAEDRLGVPTINAHPSKYDYNYWGQGTQVTVSSEPKTPKPQSGGGGSGGGGSGGGGQGVQAQLQLVESGGGLVQAGGSLRLSCAASGRTFSSYSMGWFR QAPGKEREYVAAVNSNGDSTFYADSIKGRFTVSRDAAKNTVYLQMNSLKPEDTALYYCAAVYGRYTYQSPKSYEYWGQGTQVTVSSEPKTPKPQSARQGAPVPYPDPLEPRGGGSDICLPRWGCLWED***

**VNA2-ABE**

**METDTLLLWVLLLWVPGSTGDAAQPARRARRTKLSGAPVPYPDPLEPRAAAGQGVQAQLQLVESGGGLVHPGGSLRLSCAPSASLPSTPFNPFNNMVGWYRQAPGKQREMVASIGLRINYADSVKGRFTISRDNAKNTVDLQMDSLRPEDSATYYCHIEYTHYWGKGTL VTVSSEPKTPKPQSGGGGQGVQAQLQLVESGGGLVQAGGSLRLSCAASILTYDLDYYYIGWVRQAPGKEREGVSCISSTDGATYYADSVKGRFTISRNNAKNTVYLQMNNLKPEDTAIYYCAAAPLAGRYCPASHEYGYWGQGTQVTVSSAHHSEDPSGGGGQGVQAQL QLVESGGGLVKPGGSLRLSCVVSGFTFDDYRMAWVRQAPGKELEWVSSIDSWSINTYYEDSVKGRFTISTDNAKNTLYLQMSSLKPEDTAVYYCAAEDRLGVPTINAHPSKYDYNYWGQGTQVTVSSEPKTPKPQSGGGGQGVQAQLQLVESGGGLVQVGGSLRLSCVVS GSDISGIAMGWYRQAPGKRREMVADIFSGGSTDYAGSVKGRFTISRDNAKKTSYLQMNNVKPEDTGVYYCRLYGSGDYWGQGTQVTVSSAHHSEDPSGGGGQGVQAQLQLVESGGGLVQAGGSLRLSCAASEFRAEHFAVGWFRQAPGKEREGVSCVDASGDSTAY ADSVKGRFTISRDNNKNVVYLQMDSLEPEDTGDYYCGASYFTVCAKSMRKIEYRYWGQGTQVTVSSEPKTPKPQSGGGGQGVQAQLQLVESGGGLVQAGGSLRLSCAASGRTFSSYSMGWFRQAPGKEREYVAAVNSNGDSTFYADSIKGRFTVSRDAAKNTVYLQMN SLKPEDTALYYCAAVYGRYTYQSPKSYEYWGQGTQVTVSSEPKTPKPQSARQGAPVPYPDPLEPRGGGSDICLPRWGCLWED***

**Figure S1.** Sequences of heterohexamer VNAs and their component VHHs. A. The names and amino acid sequences of the six VHH components of VNA1-ABE and VNA2-ABE are provided. B. The complete encoded amino acid sequences of both VNA1-ABE and VNA2-ABE are shown.
